# SCALE: Unsupervised Multi-Scale Domain Identification in Spatial Omics Data

**DOI:** 10.1101/2025.05.21.653987

**Authors:** Behnam Yousefi, Darius P. Schaub, Robin Khatri, Nico Kaiser, Malte Kuehl, Cedric Ly, Victor G. Puelles, Tobias B. Huber, Immo Prinz, Christian F. Krebs, Ulf Panzer, Stefan Bonn

## Abstract

Single-cell spatial transcriptomics enables precise mapping of cellular states and functional domains within their native tissue environment. These functional domains often exist at multiple spatial scales, with larger domains encompassing smaller ones, reflecting the hierarchical organization of biological systems. However, the identification of these functional domain hierarchies has been hardly explored due to the lack of appropriate computational methods. In this work, we present SCALE, an unsupervised algorithm for multi-scale domain identification in spatial transcriptomics data. SCALE combines neural graph representation learning with an entropy-based search algorithm to detect functional domains at different scales. It reaches state-of-the-art performance in single- and multi-scale domain detection on simulated and murine brain Xenium and MERFISH data, as well as patient-derived kidney tissue, highlighting its robustness and scalability across diverse tissue types and platforms. SCALE’s ease of use makes it a powerful aid for advancing our understanding of tissue organization and function in health and disease.

## Introduction

The rapidly growing field of single-cell spatial transcriptomics (ST) enables the identification of cellular states and signaling in their native microenvironments. Within a tissue, cells are often organized into distinct spatially related regions that correspond to specific anatomical structures or functional areas. These anatomical and functional domains exist across multiple scales, often in a nested structure, ranging from local cellular domains to broader anatomical domains (Fig. 1a). Identifying spatial domains across multiple scales can improve our understanding of organ function, development, and disease pathology. However, despite significant advancements in spatial molecular imaging technologies, such as MERFISH^1^ and Xenium^2^, multi-scale domain identification remains an underexplored area, and current methodologies still face substantial challenges in accurately identifying spatial domains, particularly in an unsupervised manner.

**Fig. 1.**
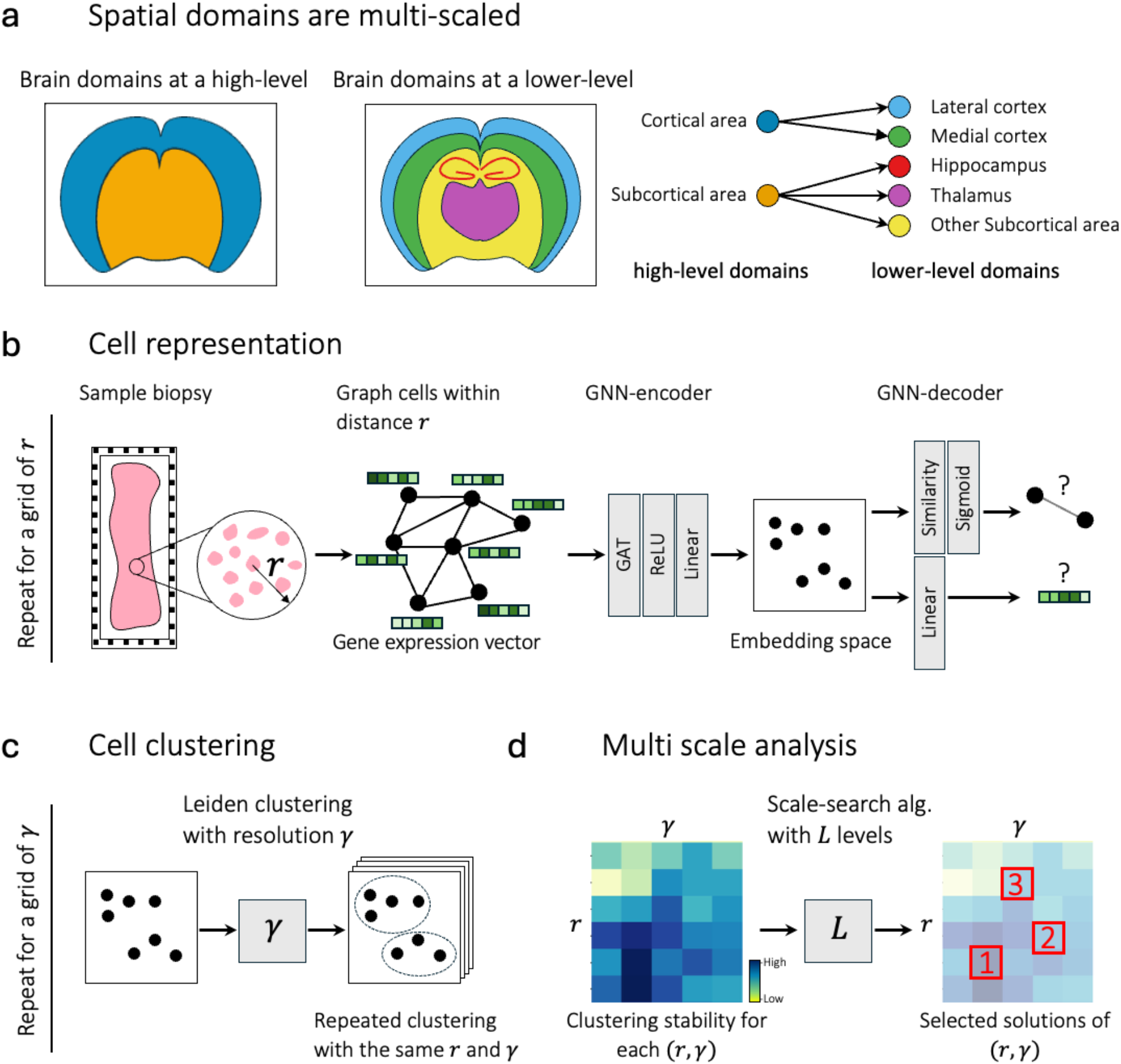
Schematic representation of spatial domain hierarchies and SCALE’s architecture. (a) Spatial domains in the brain are organized in a hierarchical, nested structure. **(b)** Given spatial transcriptomic data from a tissue biopsy, a spatial graph is constructed where cells are represented as nodes and edges represent cell-cell adjacencies within a threshold *r*. For a grid of different thresholds *r*, a graph neural network-based encoder-decoder learns cell representations by reconstructing the cellular adjacencies and gene expression. **(c)** Leiden clustering with several resolutions *γ* is used for clustering nodes in embedding space, and a cluster similarity metric measures the stability of clusters generated by each pair of *r* and *γ*. The base assumption is that functional domains should be more stable. **(d)** Our entropy-based search algorithm finds values of *r* and *γ* for a desired number of levels.

Traditional approaches in ST have often adapted clustering algorithms initially designed for single-cell RNA sequencing^3–5^, which do not take spatial information into account. Recent works have introduced spatially informed algorithms ranging from graph neural network-based methods^6–12^ to Bayesian methods^13^. Despite these advances, these methods are not designed for capturing multi-scale spatial structures. To identify domains across multiple scales, existing methods usually require manual tuning of multiple hyperparameters, while the results may highly depend on the user’s choice. Currently, the only method for the identification of hierarchical domains is NeST^14^, which searches regions of highly co-expressed genes. NeST is based on an assumption that domains in a tissue are characterized by specific gene coexpression patterns; however, this assumption may not be generalizable across different tissues, specifically where distinct domains include overlapping cell types, potentially leading to suboptimal performance.

This manuscript introduces SCALE (Spatial Clustering At multiple LEvels), a method that employs a graph neural network-based (GNN-based) encoder-decoder architecture with a bi-objective function integrating both cell transcriptomic data and spatial relationships among cells. GNN representations are subsequently subjected to a novel entropy-based search algorithm that enables the identification of optimal domains across multiple scales. SCALE is easily applicable and identifies spatial domains in simulated and real-world spatial single-cell data obtained from a variety of tissue types and technologies with best-of-breed accuracy and reliability.

## Results

### Overview of SCALE

Our algorithm is derived from three key assumptions about spatial domain organization. The coherence assumption states that cells within a spatial domain have similar gene expression in their neighborhood. The spatial continuity and scale relevance assumption asserts that biological domains form contiguous regions at specific biologically meaningful scales. The hierarchical organization premise posits that functional regions are organized in a spatially nested manner.

#### Deep learning model architecture

SCALE employs a GNN-based encoder-decoder architecture to map cells into an embedding space and subsequently utilizes a clustering algorithm to identify spatial domains. The overall framework of SCALE is illustrated in Fig. 1b-d. Having a spatially resolved sample biopsy, we first construct a spatial graph, with nodes representing cells and their gene expression profiles as node attributes. The edges represent spatial adjacency between cells defined by a spatial distance threshold *r*. In the encoder, a graph attention network (GAT)^15^ is trained to perform message passing between adjacent cells and embed them into an embedding space. The decoder is subsequently trained to reconstruct both the gene expression profiles of the cells and the cell-cell adjacency relationships (Fig. 1b). This approach ensures that, in the embedding space, cells that are spatially close and share similar gene expression neighborhoods will be positioned close to each other. Therefore, to train the model, we proposed to use a bi-objective cost function optimized along a Pareto front:

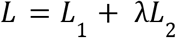

where *L*_1_ is a binary cross-entropy loss for predicting pairs of cells within a neighborhood and *L*_2_ is a mean squared error for gene expression prediction. The hyperparameter λ enables navigation across the Pareto front, balancing the trade-offs between the objectives. For a given *r*, we train our model with different values of λ and choose the one that maximizes the correlation of the GNN embedding space and spatial adjacencies, as quantified by Moran’s I (MI, Supplementary Algorithm 1). After the model is trained and cells are represented in the embedding space, we apply Leiden clustering^16^ to identify spatial domains (Fig. 1c).

#### Multi-scale domain identification

The identified domains largely depend on the choice of *r* and the Leiden resolution *γ*. Selecting these parameters is challenging for two reasons: (i) domain identification is an unsupervised task with no ground truth in real-life scenarios, and (ii) there is often no single optimal scale. In reality, however, biological tissues often contain multiple biologically meaningful spatial domains that are organized hierarchically across different scales. To address this challenge, we developed an entropy-based search algorithm that identifies a set of optimal *r* and *γ* values, enabling multi-scale domain identification (Fig. 1d). We start by generating clusters multiple times across a grid of *r* and *γ* values, then assess the stability of the resulting clusters. This approach operates on the assumption that more robust clusters are likely to be better aligned with underlying biological structures. To achieve this, we compute the adjusted rand index (ARI), *i*.*e*., a measure for evaluating the similarity between two data clusterings, for each pair of clusterings and then calculate the average ARI across all pairs, producing a stability matrix. Pairs of *r* and *γ* values associated with higher stability were retained, with each pair representing a domain clustering at a specific scale. Next, we use our entropy-based search algorithm to find spatial domains across multiple scales, optimizing for a nested structure, where lower-scale domains are contained within those at higher scales. Specifically, given an arbitrary cell in a cluster from a lower-level domain clustering, the algorithm maximizes the probability that the same cell is assigned to a single domain inside a high-level domain clustering (Supplementary Fig. 1). The mathematical details of SCALE are described in the Methods section.

### SCALE achieves state-of-the-art single-scale domain identification performance

We first assessed SCALE’s single-scale domain identification performance by comparing it to six state-of-the-art competitors: NichePCA^17^, MENDER^18^, Space-Flow^10^, SCAN-IT^6^, CellCharter^19^, Banksy^20^, and NEST^14^. Since all the existing methods, except for NeST, lack support for multi-scale domain identification, our comparison in this section is limited to single-scale domain identification. For the data, we assembled two independent spatial single-cell mouse brain datasets. Dataset 1 contains four samples measured with the MERFISH technology^1^ with 483 genes and ∼80,000 cells per sample. The ground-truth domain annotations were performed using the same automated annotation workflow as in Schaub et al. (2024)^17^, obtaining annotations for major brain regions (Level 1) and subregions (Level 2) based on the Allen mouse brain atlas^21^. For single-scale domain identification, we used Level 2 annotations as they resemble the annotations used in Dataset 2 more closely. Dataset 2 contains three mouse brain samples measured with Xenium technology^2^, featuring 248 unique genes and ∼150,000 cells per sample. The ground-truth annotations for cells and spatial domains were obtained from the original publication^22^.

The identification of spatial domains was assessed using the ground truth annotations based on different cluster similarity metrics. First, we compared SCALE with its automatic hyperparameter selection to competing methods with default parameters. However, all Leiden-based competitors still required supervised tuning of their resolution parameter to optimize for the best match to the ground truth labels, while SCALE selected it automatically in an unsupervised manner. The only constraint we imposed on SCALE was a minimum of 30 clusters in the selectable clusterings. In this setting, SCALE outperformed all competing algorithms with NichePCA being a close second (Fig. 2a and b, and Supplementary Fig. 2). On Dataset 1, SCALE achieved a median AMI of 0.67, surpassing NichePCA’s, MENDER’s, and NeST’s median AMI by 4.9, 7.6, and 211.7 percentage points, respectively. On Dataset 2, SCALE achieved a median AMI of 0.66, which is almost on par with NichePCA’s AMI of 0.67, while surpassing MENDER’s and NeST’s median AMI by 6.2 and 180.8 percentage points, respectively.

**Fig. 2.**
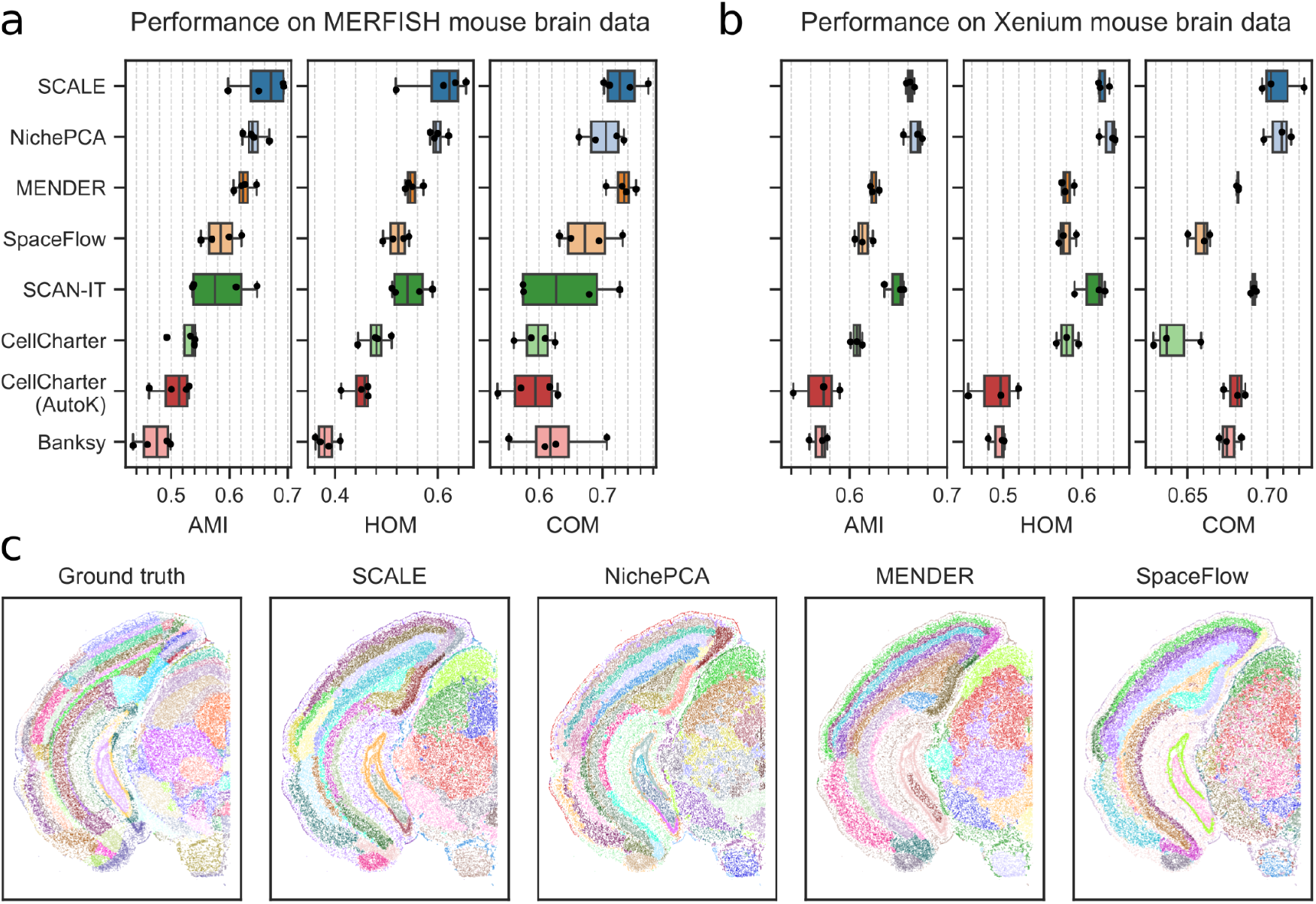
Single-scale domain detection performance comparison on Xenium and MERFISH mouse brain data. (**a**) Benchmarking performance across different methods for Xenium Dataset 1. Dots show individual sample performance, while the boxplot displays the median, the 25th and 75th percentiles, and whiskers extending 1.5 times the interquartile range. (**b**) Benchmarking performance across different methods for MERFISH Dataset 2. (**c**) Example images of the ground truth domains for the left brain hemisphere of a selected sample of the MERFISH dataset and the identified domains for SCALE, NichePCA, MENDER, and SpaceFlow.

In our next analysis, we tuned all available hyperparameters of the competing methods (e.g., the number of nearest neighbors for constructing the spatial graph) in a supervised manner to achieve the best possible single-scale domain detection performance. In this case, SCALE, although effectively unsupervised with the only constraint that candidate clusterings contain at least 30 clusters, performs on par with NichePCA and outperforms all other methods (Supplementary Fig. 3). The identified spatial domains by different methods, along with their ground truth annotation for a selected sample of Dataset 1, are shown in Fig. 2c.

In summary, SCALE provides state-of-the-art supervised and unsupervised single-scale domain detection performance, indicating that its GNN-based architecture and dual optimization function are well-suited for the task.

### SCALE can identify domains across multiple scales

While SCALE achieves state-of-the-art single-scale domain detection performance, it is specifically designed for multi-scale, unsupervised domain detection. In this section, we use one simulated and two real-world datasets to compare SCALE to NEST, the only other algorithm capable of multi-scale domain identification.

#### Multi-scale analysis in simulated data

In our simulation study, we manually crafted an image of spatial domains with predefined structures at two levels. To generate spatial single-cell data, we randomly sampled 1,000 points uniformly across a hand-crafted image representing the cell coordinates (Fig. 3a). A gene expression vector was assigned to each cell using the mouse brain data presented in Moffitt et al. (2018)^23^. At the finest scale, we considered each domain to be composed of a homogeneous mixture of randomly picked cell types. Details of the algorithm used to assign gene expression profiles to individual cells are provided in Supplementary Algorithm 2. We then applied both NeST and SCALE (with L = 2) to the simulated data to identify spatial domains (Fig. 3b, c). As can be seen, the domains identified by SCALE align almost perfectly with the ground truth labels, while NeST fails to correctly recognize the domains and leaves most cells unassigned. For a quantitative comparison of the predicted with the ground truth domains, we used AMI, HOM, and COM scores. As the results in Fig. 3d show, SCALE reliably identifies the ground truth domains at different scales, while NeST performs poorly. For high-level domain identification, SCALE achieves a median AMI score of 0.85, surpassing NeST by 158.4 percentage points. In the low-level case, SCALE reaches an AMI score of 0.75, again outperforming NeST by 42.5 percentage points.

**Fig. 3.**
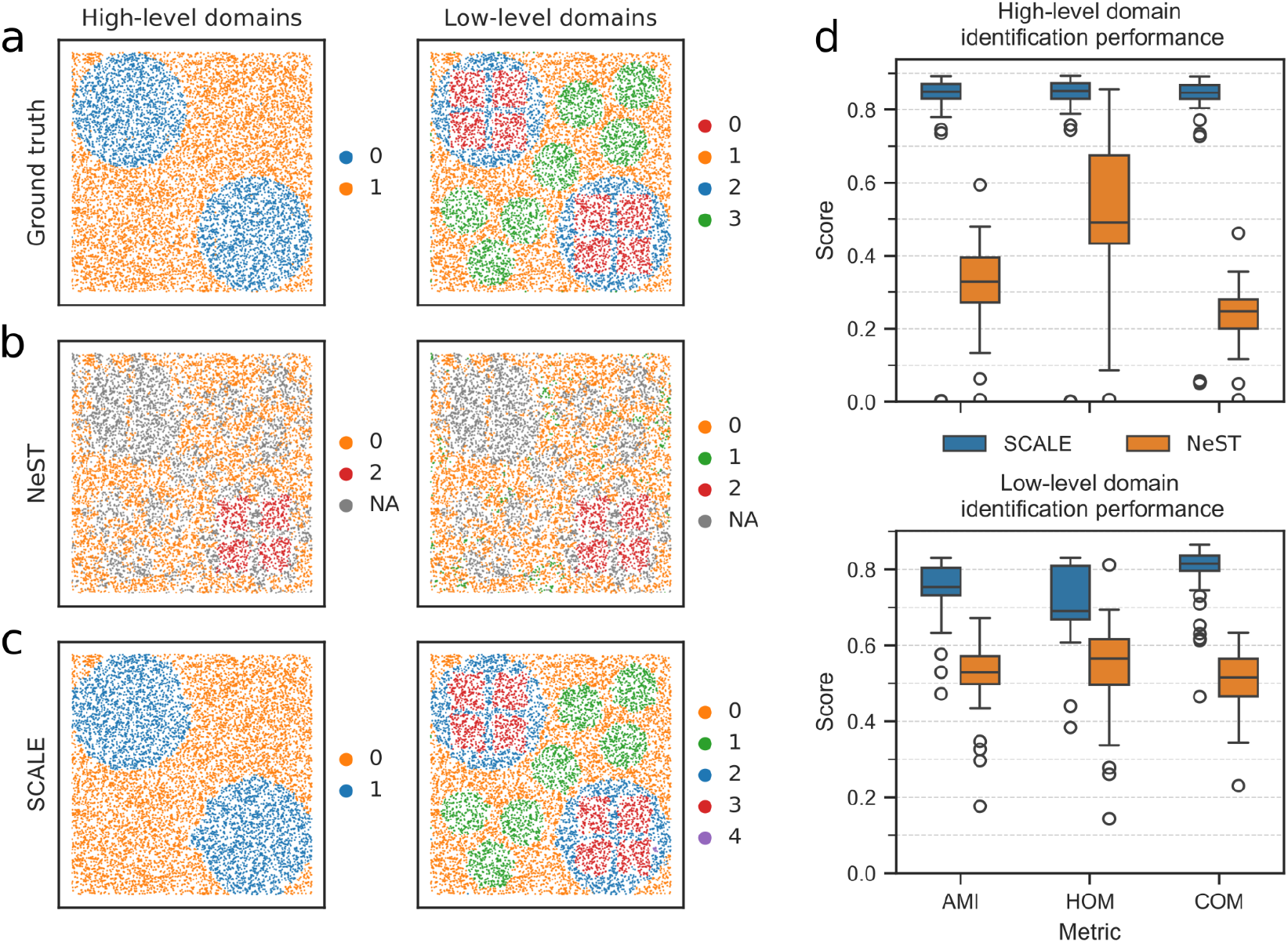
Multi-scale domain identification results on simulated data. (**a**) Ground truth high-level and low-level domain annotations, (**b)** identified domains by NeST, and (**c)** SCALE for a representative sample from our 50 simulated samples. Cells not assigned to any domain by NeST are labeled as “NA”. **(d)** Performance comparison between SCALE and NeST across all 50 simulated samples at high- and low-level scales in terms of AMI, HOM, and COM scores. On one sample, NeST and SCALE assigned all cells to a single domain, leading to near-zero scores.

#### Multi-scale analysis of mouse brain data

As an example of real-world data, we used our Dataset 1 considering both the Level 1 (high-level) and Level 2 (low-level) domains for the ground truth annotations (Fig. 4a). We used NeST and SCALE (with L = 2) to identify domains across two scales and the results are shown in Fig. 4b and 4c. NeST only detected a single scale with many cells not assigned to any cluster. Moreover, it failed to correctly recognize certain domains, in particular hippocampal subfields. SCALE, on the other hand, showed promising results in identifying multiple scales in the mouse brain data. In particular, as the high-level domains, SCALE identified the brain cortex, hippocampus, and thalamus, which are functionally important areas the brain. Each of these domains was then divided into lower-level scales such as cortical layers and hippocampal subfields. Figure 4d shows the similarity of the identified domains with the ground truth annotations at high- and low-level scales. Similar to the above analysis, we used AMI, HOM, and COM scores as the clustering metrics. SCALE demonstrates notably higher similarity to the ground-truth labels at both scales compared to NeST. Specifically, it achieves a median AMI score of 0.53 for high-level domain detection, exceeding NeST by 104.0 percentage points. For low-level domains, SCALE reaches a median AMI score of 0.62, again outperforming NeST by 191.1 percentage points.

**Fig. 4.**
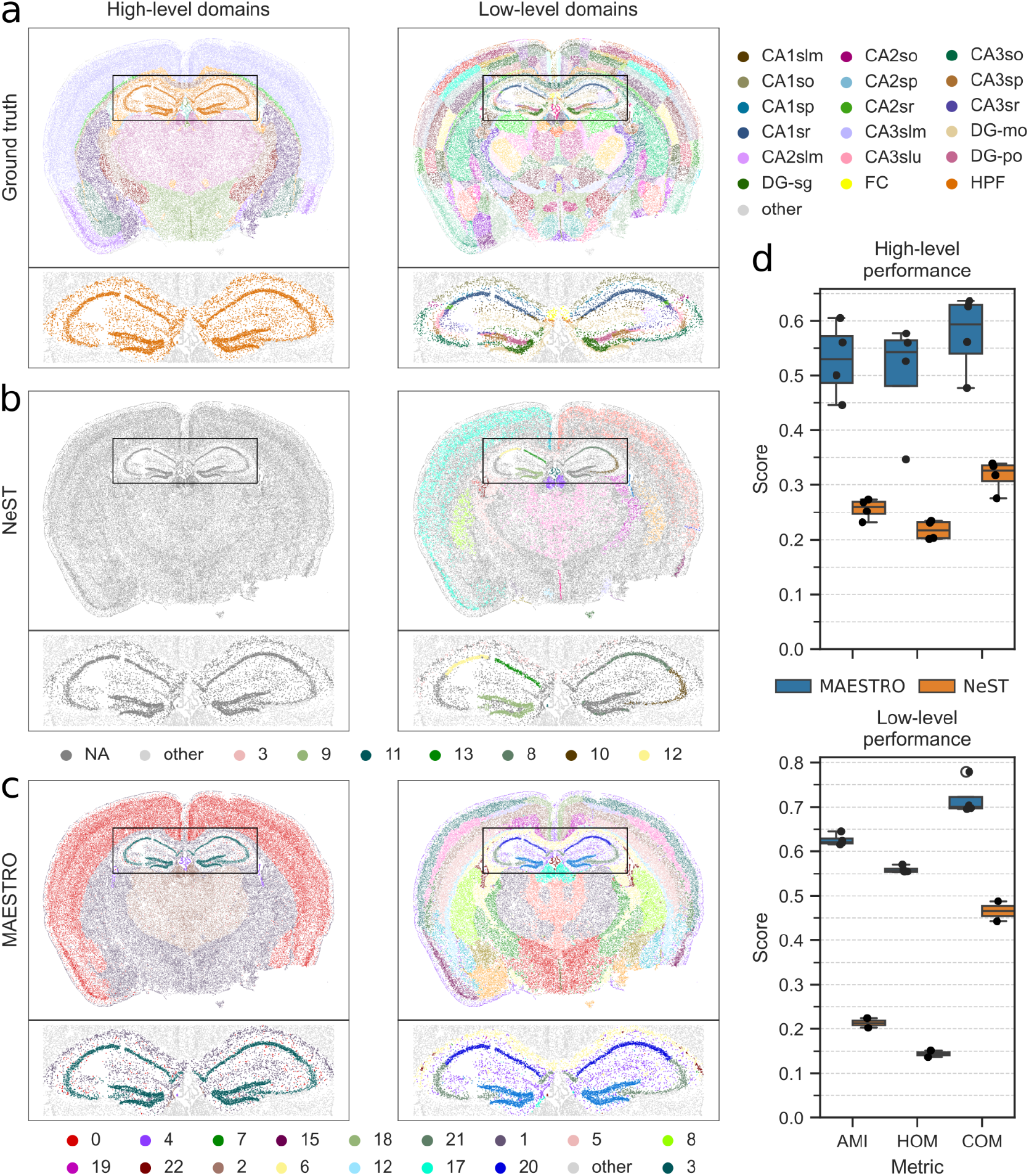
Multi-scale domain identification results on MERFISH mouse brain data. **(a)** Ground truth domain annotations for a selected sample of Dataset 1 were obtained at two levels based on the Allen mouse brain atlas. **(b)** domains identified by NeST, and **(c)** domains identified by SCALE at each scale. The inlets show the domain clusters inside the hippocampal domain; all other cells are grayed out. Cells not assigned to any domain by NeST are labeled as “NA”. Since NeST does not allow the detection of a predefined number of domain levels, it only outputs results for one level on this sample. **(d**) The similarity of identified domains with ground truth annotations at high- and low-level scales as measured by AMI, HOM, and COM scores. NeST identified domains at two scales only for two of the four samples.

#### Multi-scale analysis in human kidney data

We further demonstrate SCALE’s ability to delineate distinct domains in patient-derived kidney biopsies.. For this, we used the data from Sultana et al.^24^, which consists of gene expression information for a custom gene panel of 480 genes at subcellular resolution measured on tissue sections of the human kidney using the Xenium technology. We employed NeST and SCALE (L=2) to identify domains on a control sample (healthy regions of a tumor nephrectomy specimen) shown in Supplementary Fig. 5. Then, we used genetic biomarkers to identify different kidney compartments, including glomeruli, proximal convoluted tube, distal convoluted tube, loop of Henle, and vasculature^25,26^ (Supplementary Fig. 4). Fig. 5 shows domains annotated for the higher- and lower-level scales, respectively. NeST only detected a single scale with 58 clusters and a large region of the tissue was not assigned to any cluster, here labeled as “NA” (Supplementary Fig. 5a), which did not align well with the marker genes related to the kidney compartments (Supplementary Fig. 6a). Nevertheless, in the results obtained by SCALE (Fig. 5b), the higher-level scale distinguishes the glomeruli domain from the tubulointerstitial regions; the lower-level scale further identifies finer domains, including proximal convoluted tube, distal convoluted tube, loop of Henle, and vasculature (Supplementary Fig. 6b,c). In addition, we assessed the performance of our method in the recognition of glomerular regions in comparison with expert manual annotations (Supplementary Fig. 7). Our method demonstrated a sensitivity of 100% and specificity of 88%, showing the capability of SCALE to reliably identify relevant domains in human biopsy data.

**Fig. 5.**
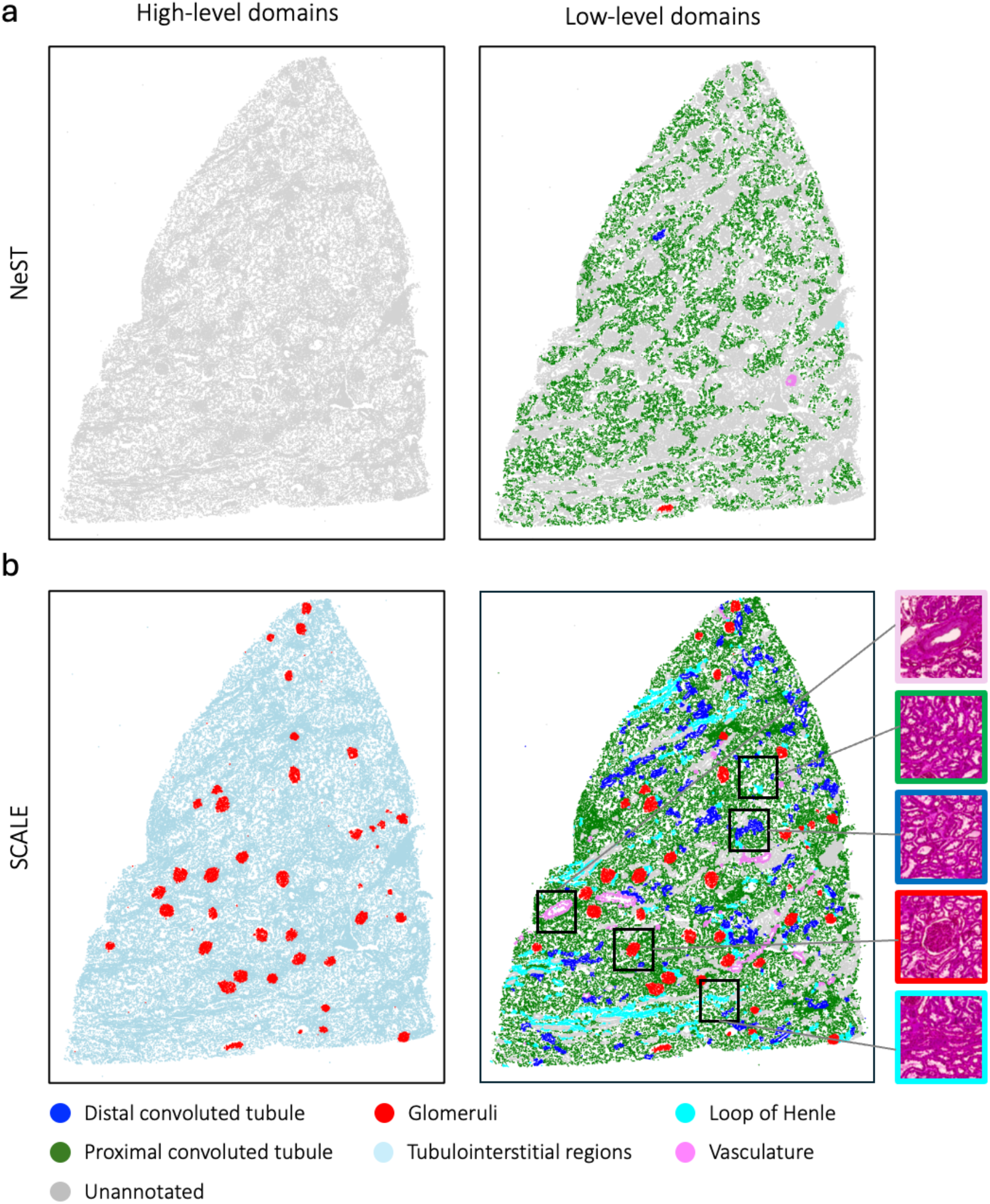
Multi-scale domain identification by SCALE and NEST on a human kidney biopsy. All annotations are based on marker genes (e.g., Glomeruli in red). Domains not assigned to any kidney compartments are denoted as ‘Unannotated’. **(a)** Domains identified by NeST, and **(b)** by SCALE at high-level and low-level scales. H&E staining images are provided to showcase kidney compartments. While SCALE reliably captures a high-level representation of glomerular and extra-glomerular regions, NEST fails to find a high-level representation.

## Discussion

Understanding tissue organization requires the identification of spatial domains and capturing of their hierarchical structure, as biological processes operate across a range of spatial scales — from small, localized niches to broader tissue-level patterns. This hierarchical organization of functional domains is essential for organ function and unsupervised charting of these intricate relationships has to date hardly been investigated. Current domain detection algorithms mostly lack the ability to flexibly detect domains at multiple scales without manual intervention, potentially overlooking important layers of biological organization. In this manuscript, we present SCALE, a deep learning framework designed for the identification of multi-scale spatial domains in single-cell spatial transcriptomics data. Leveraging a bi-objective training scheme that anchors both transcriptomic similarity and spatial proximity, SCALE learns an embedding space to simultaneously reflect the gene expression neighborhoods of cells and their spatial cellular adjacencies. In addition, SCALE introduces an innovative entropy-based search algorithm that automatically adapts its key parameters *r* and *γ*, enabling robust domain detection across scales with minimal manual calibration. Together, these advances position SCALE as a versatile and scalable tool for decoding tissue organization from spatial omics data.

To the best of our knowledge, NeST is the only method specifically designed to identify spatial domains across multiple scales. However, in all of our analyses, NeST resulted in suboptimal domain identification performance. In contrast, SCALE showed promising results when tested for multi-scale domain identification based on simulated data, mouse brain tissue, and kidney tissue with both MERFISH and Xenium technologies. These results prove SCALE’s versatility across different tissues, ST technologies, and organisms. In the mouse brain tissue, SCALE identified the primary anatomical domains as high-level domains, while simultaneously detecting finer-scale organization, such as cortical layers and hippocampal compartments at a lower level. In the kidney tissue, SCALE annotated glomerular and tubulointerstitial regions as the two high-level domains. At the lower level, SCALE delineated proximal convoluted tube, distal convoluted tube, loop of Henle, and vasculature as distinct compartments.

We benchmarked SCALE against the state-of-the-art in a single-scale domain identification setting and showed its accuracy and robustness in identifying spatial domains for two independent mouse brain datasets. In particular, we conducted two experiments, both with SCALE optimized in an unsupervised manner. In the first experiment, all methods were run with default parameters, tuning only the Leiden resolution, where SCALE outperformed the competing algorithms. In the second experiment, SCALE was compared against methods optimized using a supervised approach. Although it showed slightly lower performance than NichePCA, it still outperformed or matched all other algorithms. These results demonstrate SCALE’s robustness and adaptability, as it maintains superior performance even when optimized in an unsupervised manner, highlighting its potential for real-world applications where supervised optimization is not feasible.

Despite the superior performance and applicability of SCALE, it does have certain limitations. While its automatic hyperparameter tuning enables effective multi-scale domain identification, setting it apart from other methods, it can significantly increase computational time. Users, however, have the flexibility to bypass our hyperparameter tuning and input their own parameters. Future work could focus on enhancing the efficiency of this process by integrating faster search algorithms, such as evolutionary search methods, to improve the scalability and speed of SCALE further. We also acknowledge that some low-level domains, such as vasculature, may span multiple high-level domains, making it difficult for SCALE to capture them as coherent units. This hierarchical overlap may lead to the exclusion of biologically meaningful structures that are not confined to a single higher-level domain. Future work will be necessary to better handle these corner cases.

Overall, SCALE represents a significant advancement in unsupervised, multi-scale spatial domain identification by leveraging graph neural networks and information theory to uncover nested functional structures within spatial transcriptomic data. Unlike existing methods, SCALE reliably detects hierarchical domains across diverse tissue types and technologies, demonstrating superior performance in both simulated and real-world datasets. Its ability to generalize across different spatial omics platforms and tissue types underscores its adaptability and real-world applicability. SCALE provides a powerful framework for advancing our understanding of tissue organization in both health and disease, with potential implications for precision medicine and clinical decision-making.

## Online Methods

### SCALE Description

Given a set of *n* cells *C* = {*c*_1_, …, *c*_*n*_} in a tissue, each with *s* dimensional spatial coordinates *y*_*i*_ ∈ *R*^*s*^ and *g* dimensional gene expression profiles *x*_*i*_ ∈ *R*^*g*^, we developed a SCALE to identify spatial domains 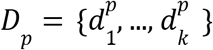 at scale *p*, where the number of domains *k* is unknown a priori. Each domain 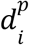 is a subset of *C* such that *d*_*i*_ ∩ *d*_*j*_ = ∅; ∀ *i*≠*j*.

### Assumptions

Our approach is derived from three minimal assumptions about the nature of spatial domains in biological tissues:

1. Local coherence: cells in a spatial domain have similar gene expression profiles in their neighborhood.
2. Spatial continuity and scale relevance: biological domains form spatially contiguous regions, with their structure being most pronounced at specific biologically meaningful scales.
3. Hierarchical organization: functional domains generally exhibit a spatially nested structure.

To satisfy our assumptions, we first defined a spatial graph and then used GNN-based representation learning to embed cells into a vector space. We introduced a bi-objective function to integrate both the gene expression similarity and spatial proximity representation while training the model. We then presented a scale-tuning algorithm to identify domains at multiple nested tissue scales.

### Representing spatial proximity as a spatial graph

Let *G* = (*G V, E*_*r*_, *X*) be a graph with the sets of nodes (vertices) *V* = {*v*_*i*_}, edges 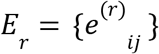, and node features *X* ∈ *R*^*n*×*g*^. To define a spatial graph, we let *V* = *C* represent the cells as nodes and *X* denote the expression profiles of cells. *E*_*r*_ represents spatial proximity within the distance threshold *r*. In particular, the existence of an edge *e*^*(r)*^_*ij*_ between cells *c*_*i*_ and *c*_*j*_ determined by:

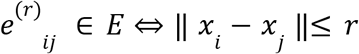

### GNN-based representation learning

While several architectures can be used to incorporate our assumptions, a GNN-based model is naturally suitable as it inherently takes into account spatial proximity as a graph and is flexible to ensure local coherence. We defined the model architecture below:

Let *h* be the dimension of the learned embedding space *Z* ∈ *R*^*n*×*h*^. The network architecture consists of an encoder for graph representation learning, a link prediction decoder, and a gene expression decoder. The encoder learns a mapping function *f*_θ_ that maps *G*_*r*_ into the embedding space *Z*_*r*_,

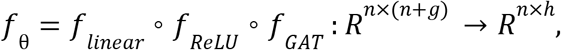

The link prediction decoder maps the embeddings of a pair of cells onto a scalar value as a probability representing the existence of an edge,

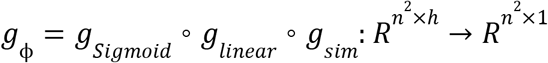

Where *g*_*sim*_ (*i, j*) =− ||*f*_θ_ (*i*) − *f*_θ_ (*j*)||^2^ ∀*i, j s. d. e*_*ij*_ ∈ *E*. Finally, the gene expression decoder maps the embedding of each cell onto a reconstructed gene expression vector,

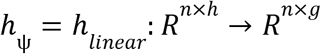

### Objective function

We defined a bi-objective function that incorporates both decoders of spatial and molecular objectives as:

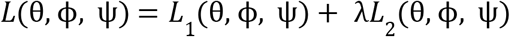

where:

- *L*_1_ = *BCE*(*g*_*ϕ*_(*f* _θ_ (*G*_*r*_)), *A*) is the binary cross-entropy (BCE) loss for link prediction, encouraging the model to capture local spatial structure.
- *L*_2_ = *MSE*(*h*_*Ψ*_ (*f*_θ_ (*G*_*r*_)), *Y*) is the mean squared error (MSE) for gene expression prediction, ensuring the embeddings capture global gene expression patterns.

### Pareto Optimization for λ

To find optimal trade-offs between spatial and molecular objectives, we trained multiple models with different λ values: 0.000001, 0.000005, 0.00001, 0.00005, 0.0001, 0.0005, 0.001, 0.005,

0.01, 0.05, 0.1, 0.5, 1, 5, 10, 50, 100, 500, 1000. For each value of λ, we calculated the correlation between the GNN embedding space and spatial adjacencies based on Moran’s I (MI) (using the *morans_i* function from the Scanpy v1.10.1 Python package). To determine the optimal λ, we first identified the saturation point on the *MI*-λ curve. If no saturation point was observed, we selected the λ value corresponding to the maximum observed MI. The algorithm is presented in Supplementary Algorithm 1.

### Domain identification by clustering

For a given radius *r*, we constructed *G*_*r*_ and computed the embeddings *Z*_*r*_ = *f*_θ_ (*G*_*r*_). Next, we applied a hard clustering method on *Z*_*r*_ to identify spatial domains satisfying *d*_*i*_ ∩ *d*_*j*_ = ∅; ∀ *i*≠*j*. Here, we used the Leiden clustering algorithm as implemented in the Scanpy Python package. An important hyperparameter of Leiden is the resolution *γ* that controls the number of clusters. Hence, the final clustering *D* is a function of (*r, γ*).

### Multi-scale domain identification by optimizing *r* and *γ*

Each value of (*r, γ*) can represent a potential solution at a different scale. To identify an optimized set of (*r, γ*) values, we implemented a two-step scale-tuning algorithm approach designed to ensure: (i) cluster stability, and (ii) a nested structure of the clusters from different scales. These two steps are described in the following:

The final clustering *D*(*r, γ*), to some degree, can depend on algorithmic initializations. We denote this variation using an index *i* as *D*^(*i*)^(*r, γ*) (*r* between 15 *μm* and 55 *μm* and *γ* between 0.01 and 1.2, see Supplementary Table 1). Among the clusterings obtained from different scales (*r, γ*), we selected those with larger stability, i.e., being less sensitive to random initializations^27^. To define the stability of clusters generated based on (*r, γ*) we used a measure of cluster similarity, in particular ARI, as

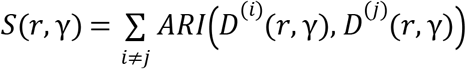

High *S*(*r, γ*) indicates stable domain identification, while low *S*(*r, γ*) suggests the scale (*r, γ*) may not be appropriate for domain identification. Calculating the cluster stability allows us to determine a set of the most stable clusterings for the data. We selected the top k (15%) most stable solutions for subsequent analysis, assuming all biologically meaningful domain scales are contained in this set. Next, we considered that domains identified at lower-level scales will generally be nested within those identified at higher-level scales. Accordingly, we introduced a novel entropy-search algorithm to find sets of (*r, γ*) for a desired number of scales, optimizing for a nested structure where lower-scale domains align with those at higher scales. The algorithm searches for a pair of domain clusterings 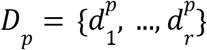 and 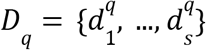 at scales *p* and *q* that best adhere to the nestedness assumption. We achieve this by maximizing the following entropy score:

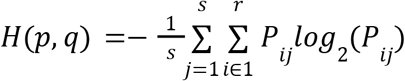

where *P*_*ij*_ is the probability that any cell *x* inside the lower-level cluster 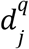 is also contained in the higher-level cluster 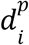 defined as

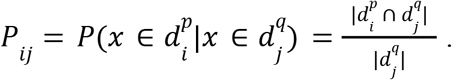

This approach naturally extends to *l* scales by iterating over all possible *l*-tuples of (*r, γ*), ordered by the number of clusters. For each *l*-tuple, we compute the entropy score *H* across all *l* − 1 cluster pairs it contains. The *l*-tuple with the lowest average entropy score is then selected as the final multi-scale domain solution. The user can also specify the minimum number of clusters to be added at each scale (we set this to 20 for the mouse brain data, and 5 for the kidney data).

### Domain annotation based on marker genes

After domain identification, we annotated the kidney compartments using the compartment-specific marker gene sets (Supplementary Fig. 4). For each gene set, we calculated a score by computing the average expression of its constituent genes. These scores were averaged across all cells in a domain to obtain domain-specific kidney compartment scores. Finally, we assigned the kidney compartment with the highest score to a domain (Supplementary Fig. 6).

## Data

### Dataset 1

This publicly available dataset was generated using MERFISH technology by the manufacturer Vizgen. It contains measurements for 483 genes, with cell segmentation performed using Vizgen’s default pipeline. We applied the same domain annotation workflow as Schaub et al. (2024)^17^ to the four most symmetric samples. Each of these four samples contains approximately 80,000 cells. Links to the original data are provided in Supplementary Table 2.

### Dataset 2

This publicly available dataset comprises three mouse brain samples measured using Xenium technology by 10x Genomics. Each sample includes approximately 150,000 cells with expression data for 248 unique genes. Cells were segmented using the proprietary pipeline from 10x Genomics. The dataset was recently annotated by Bhuva et al. (2024)^22^, who mapped spatial domains at the transcript level based on the Allen Brain Reference Atlas^21^. We followed the same preprocessing steps as in Schaub et al. (2024)^17^ and excluded dissociated cells by constructing a cell graph using a 60-micron distance threshold and retaining only cells within the largest connected component. The data was downloaded from the resource provided by Bhuva et al. (2024)^22^, and the corresponding links are provided in Supplementary Table 2.

For all datasets, we first removed cells containing fewer than 10 transcripts and genes expressed in fewer than 5 cells. Then, we normalized the raw spatial transcriptomics data to sum to their medians and applied a log-transformation, respectively, using the *normalize_total* and *log1p* functions of the Scanpy (v1.10.1) Python package^3^.

### Dataset 3

Sultana et al.^24^ generated spatially resolved gene expression data from 64 kidney biopsies. Gene expression was profiled at subcellular resolution using the 10x Xenium platform, employing a custom panel of 480 genes on tissue sections.

### Evaluation metrics

To evaluate the performance of SCALE in recognizing the “ground-truth” labels and compare it with other methods, we used the following metrics. All of these metrics were implemented via the scitkit-learn (v1.3.0) Python package^28^.

### Adjusted mutual information score (AMI)

AMI quantifies the similarity between two clusterings *U* and *V* (i.e. in our case, the model-generated clusters and the ground truth) based on their mutual information *MI*(*U, V*) while accounting for the possibility that some agreement between clusterings could happen randomly. AMI is defined as

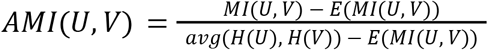

where *MI*(), *E*() and *avg*() represent the mutual information, expectation, and average, respectively. The score is normalized, with a value of 1 indicating perfect clustering correspondence, a value of 0 indicating that the similarity is no better than random, and negative values showing worse than random correspondence.

### Homogeneity score (HOM)

HOM is a score evaluating the purity of clusters with respect to the ground-truth labels. It measures whether each cluster primarily contains data points belonging to a single class defined as

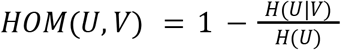

where *H*() is the Shannon entropy. A HOM of 1 indicates perfect homogeneity, meaning each cluster is entirely composed of a single class, while a score of 0 indicates poor homogeneity.

### Completeness score (COM)

COM is a score evaluating the extent to which all data points belonging to a given ground-truth label are assigned to the same cluster recognized by the models defined as

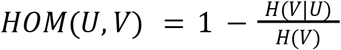

A COM of 1 indicates perfect completeness, meaning all members of each class are grouped within a single cluster, while a score of 0 suggests that members of the same class are distributed across multiple clusters.

### Adjusted rand index (ARI)

ARI is a measure used to evaluate the similarity between two data clusterings by considering both the agreements and disagreements between cluster pairs and counting pairs that are assigned in the same or different clusters in the predicted and ground-truth clusterings. ARI is defined as

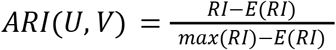

where *RI* is the rand score defined as

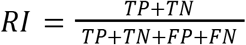

where:

- TP is the number of pairs that are in the same cluster in both U and V (true positives).
- TN is the number of pairs that are in different clusters in both U and V (true negatives).
- FP is the number of pairs that are in the same cluster in U but in different clusters in V (false positives).
- FN is the number of pairs that are in different clusters in U but in the same cluster in V (false negatives).

An ARI of 1 indicates perfect clustering correspondence, and 0 indicates random labeling.

### Benchmarking workflow

We used the ground truth annotations for Datasets 1 and 2 to quantitatively compare the performance of SCALE with the following existing spatial domain identification methods: MENDER^18^, Banksy^20^, CellCharter^19^, SCAN-IT^6^, SpaceFlow^10^. We additionally attempted to compare our method against STAGATE^8^, BASS^13^, SpatialPCA^29^, and SpaGCN^7^, but we were not able to execute them on either Dataset within a memory budget of 128 GB.

For SCALE, we applied our automatic cluster-stability-based search procedure per sample to determine the distance cutoff for spatial graph construction and the resolution for Leiden clustering (with the only restriction that the number of clusters should be at least 30). No other method allows for this kind of unsupervised parameter selection. CellCharter offers a similar procedure, called AutoK, but it only determines the number of clusters. For their algorithm, we limited the number of clusters to be between 20 and 60. To make the results for the existing methods comparable, we selected the default distance cutoff for graph construction (or similar parameters, depending on the method) while choosing the cluster resolution (or number of clusters) in a supervised fashion per sample, i.e., we selected the resolution resulting in the best AMI score. Note that this comparison favors the competing methods over SCALE.

Moreover, we compared SCALE against competing methods with tuned hyperparameters, i.e., not only the resolution (or number of clusters) but also another method-specific parameter was selected to maximize the AMI score. The parameter space for each method is provided in Supplementary Table 3.

## Computational resources

Computations were performed on an AMD EPYC 7742 central processing unit (2.25 GHz, 512 MB L3 cache, 64 cores, 128 GB RAM) and an NVIDIA A100 Tensor Core GPU with 40 GB VRAM.

## Data availability

All the raw data can be downloaded from the links provided in Supplementary Table 2. The workflows for further processing and domain annotation are described in the Methods section.

## Code availability

The code for SCALE is available at https://github.com/imsb-uke/scale. All scripts to reproduce the benchmarking and analysis are available at https://github.com/imsb-uke/scale-analysis.

## Acknowledgments

We would like to thank the members of the Institute of Medical Systems Bioinformatics for their feedback and Sven Heins and Vadim Ustinov for IT support. This study was supported by grants from the Deutsche Forschungsgemeinschaft (DFG) to UP (SFB 1192 A1 and C3), CFK (SFB 1192 A5 and C3; KR 3483/3-1), SB (SFB 1192 A2, B8, and C3), RK (FOR5068 P9), CL (SFB 1286 Z2), DPS (SFB 1192 A1). BY was supported by the Federal Ministry of Education and Research (BMBF) as part of the German Center for Child and Adolescent Health (DZKJ). NK was supported by DFG SFB1192 B8 and CDL FLIGHT of the University of Hamburg.

## Author contributions

Conceptualization and methodology: BY, DPS, and SB. Formal analysis: BY, DPS, RK, CL, MK, and NK. Writing original draft: BY, DSP, RK, CL. Reviewing and editing of the manuscript: BY, DSP, RK, CL, MK, NK, and SB. Visualization: BY, DSP, RK, CL, and NK. Supervision: VGP, TBH, CFK, UP, and SB. Funding acquisition: VGP, TBH, CFK, UP, and SB.

