## supplementary materials for "SCALE: Unsupervised Multi-Scale Domain Identification in Spatial Omics Data"

**Supplementary Algorithm 1.** Algorithm for the optimization of λ.

**Input:** a set of $R = \{r_1, ..., r_n\}$, a set of $\Lambda = \{\lambda_1, ..., \lambda_m\}$ and GNN embeddings $H_\lambda^{(r)}$ trained for each λ and $r$, a KNN neighbourhood graph ($k = 4$) $W = \{w_{ij}\}|_{i\neq j};\ w_{ij} \in \{0, 1\}$

**Output:** a set of optimum λ per $r$ $\Lambda^* = \{\lambda^*[r_1], ..., \lambda^*[r_n]\}$

1. for $r$ in $R$
2. for λ in $\Lambda^{(r)}$
3. $m[\lambda]$ = Moran's I (W, $H_\lambda^{(r)}$)
4. s[λ] = a sigmoid curve fitted on $m[\lambda]$
5. $\lambda_{saturation} = arg.min_\lambda \{\frac{d^2}{d\lambda^2} s[\lambda]\}$
6. $\lambda_{peak} = arg.max_\lambda \{s[\lambda]\}$
7. if $m(\lambda_{peak}) - m(\lambda_{saturation}) < 0.05 . m(\lambda_{peak})$
8. $\lambda^*[r] = \lambda_{saturation}$ # pick the saturation point
9. else
10. $\lambda^*[r] = \lambda_{peak}$ # pick the max value as no saturation is observed

**Supplementary Algorithm 2.** Algorithm for the assignment of gene expression values to cells in the simulated data

**Input:** a reference gene expression $X$ (*cell* × *expression*) along with the vector of corresponding cell types $T$. A set of cells $C$ with their domains $D$ at the finest scale. Noise probability $p_{noise}$

**Output:** gene expression matrix $Y$ for cells in $C$

1. for $d$ in $D$
2. $type_1[d], type_2[d]$ = randomly choose two cell types without replacement $\subset T$
3. for $i$ in $C$
4. $d = D[i]$
5. $v$ = random variable $\sim uniform(0, 1)$
6. if $v > p_{noise}$
7. $t$ = randomly choose a cell type $\in \{type_1[d], type_2[d]\}$
8. else
9. $t$ = randomly choose a cell type $\notin \{type_1[d], type_2[d]\}$
10. $cell$ = randomly select a cell of type $t$ from $T$
11. $Y[i, :]$ = m $X[cell, :]$

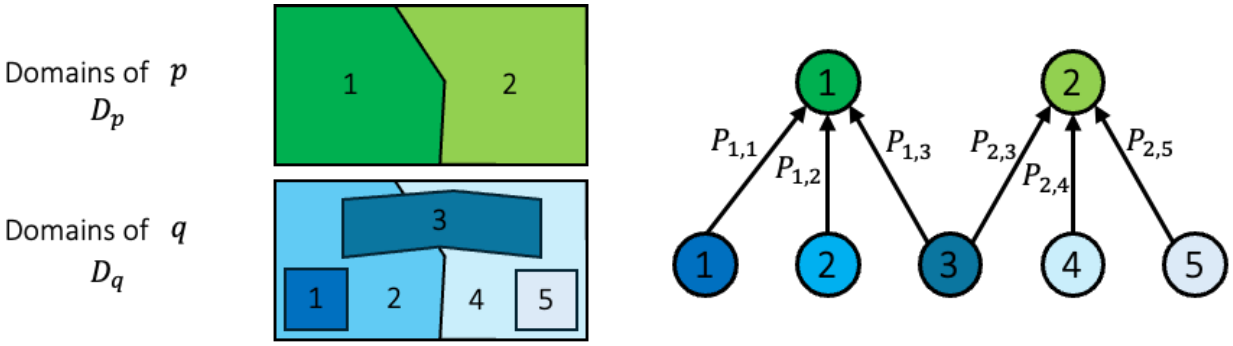


**Supplementary Fig. 1.** An example representing the entropy-based search algorithm. **(a)** sets of spatial clusters $D_p = \{d_1^p, d_2^p\}$ (shown in different shades of green) and $D_q = \{d_1^q, ..., d_5^q\}$ (shown in different shades of blue) corresponding to two different solutions $p$ and $q$. **(b)** Probability values $P_{ij} = P(x \in d_i^p | x \in d_j^q)$ can be represented as a tree. In this example, Cluster 3 at $q$ is shared between the two clusters at $p$ disrupting the nested structure and leading to an undesirable solution.

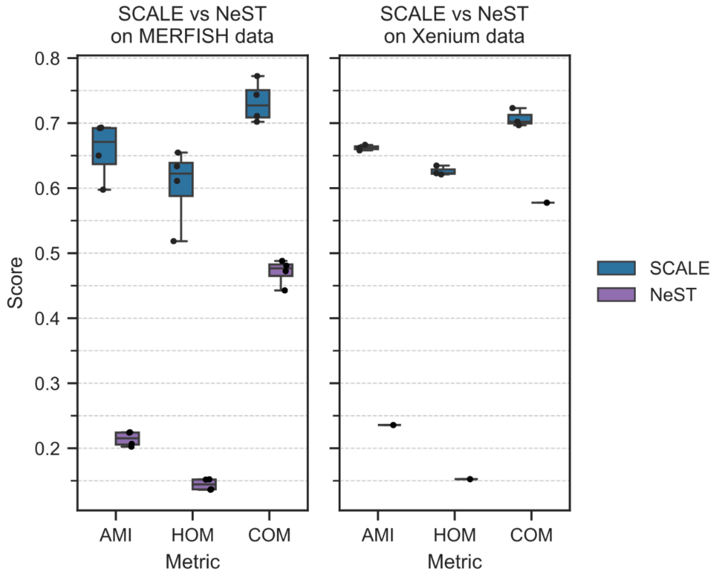


**Supplementary Fig. 2.** Performance comparison between SCALE and NeST on the MERFISH (left) and Xenium (right) mouse brain datasets regarding AMI, HOM, and COM scores. Dots show individual sample performance, while the boxplot displays the median, the 25th and 75th percentiles, and whiskers extending 1.5 times the interquartile range. For the Xenium dataset, NeST only identified domains for one sample (for different tested parameter sets).

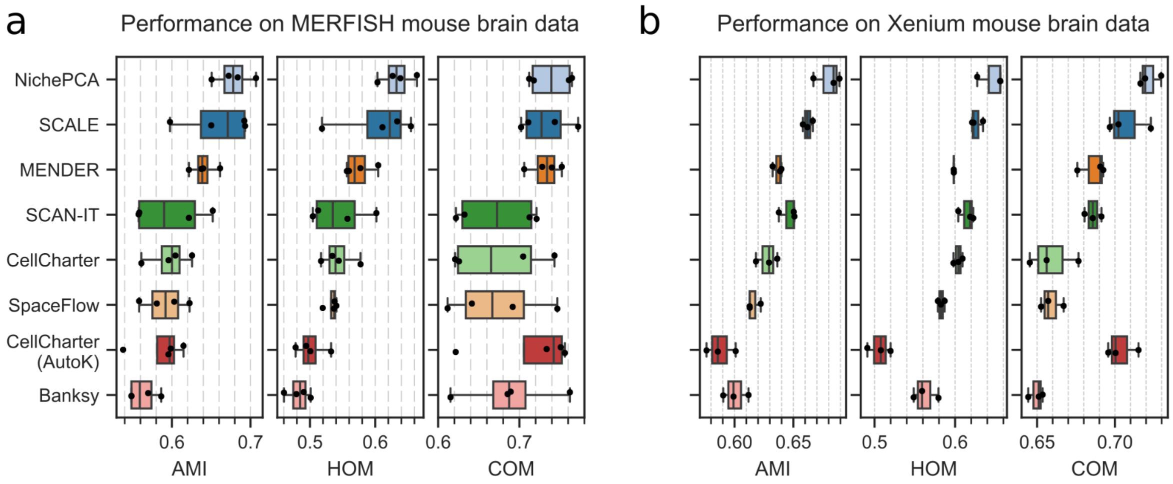


**Supplementary Fig. 3.** Spatial domain identification performance for SCALE compared against different methods with supervised hyperparameter selection on Dataset 1 **(a)** and Dataset 2 **(b)** in terms of AMI, HOM, and COM scores. Dots show individual sample performance, while the boxplot displays the median, the 25th and 75th percentiles, and whiskers extending 1.5 times the interquartile range. All results are ordered according to the average AMI score on Dataset 1.

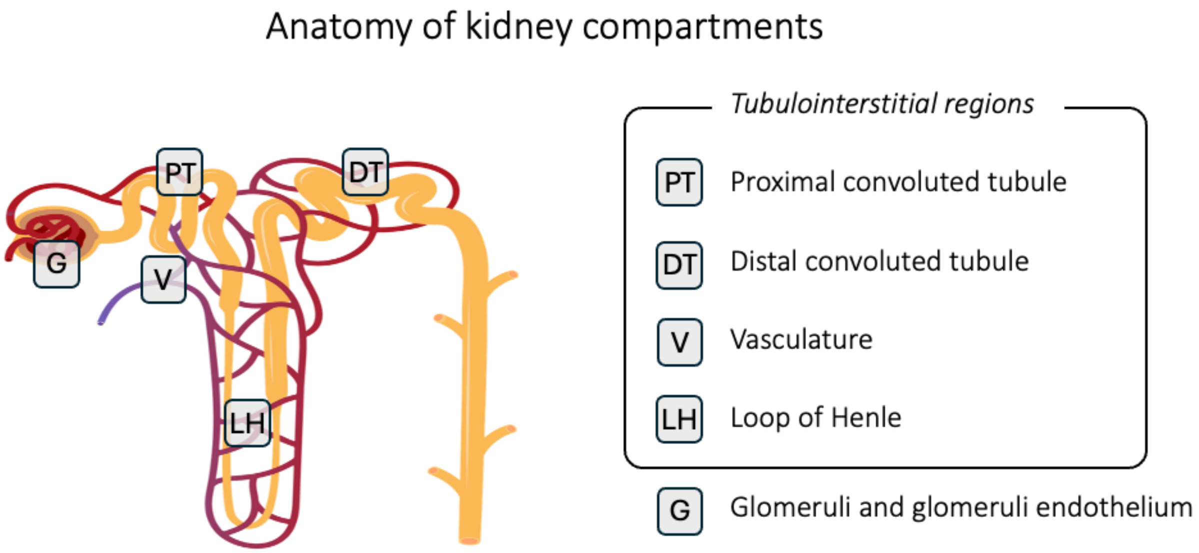


| **Domains** | **Marker genes** |
|---|---|
| Glomeruli | NPHS1, NPHS2, SYNPO, CDKN1C, WT1, FOXC2, MAFB, EFNB2, FOXL1, CD2AP, PLCE1, MYH9 |
| Glomeruli endothelium | PLAT, EMCN, TSPAN7, MAPT, KDR, SMAD6, EHD3, FLT1, KDR, BMX |
| Vasculature | NRP1, CDH5, ELN, SMAD6, LPL, FBLN2, EDN1, FBLN5, KLF4, CAS6 |
| Mesangium | SERPINE2, DES, TAGLN |
| Proximal convoluted tubule | SLC34A1, LRP2, HXYD2, HRSP12, ACSM1, ACSM2, ATP11A, CPT1A, NOTCH2, VCAM1, SLC13A3, SLC1A1, SLC5A2, SLC5A12, SLC6A19, ADRA1A |
| Distal convoluted tubule | PVALB, SLC12A3, CALB1, SLC8A1, KLK1WNK1, FXYD2, TRPM7 |

**Supplementary Fig. 4.** The anatomy of kidney compartments with the list of marker genes for each compartment. We used this gene list to annotate spatial clusters.

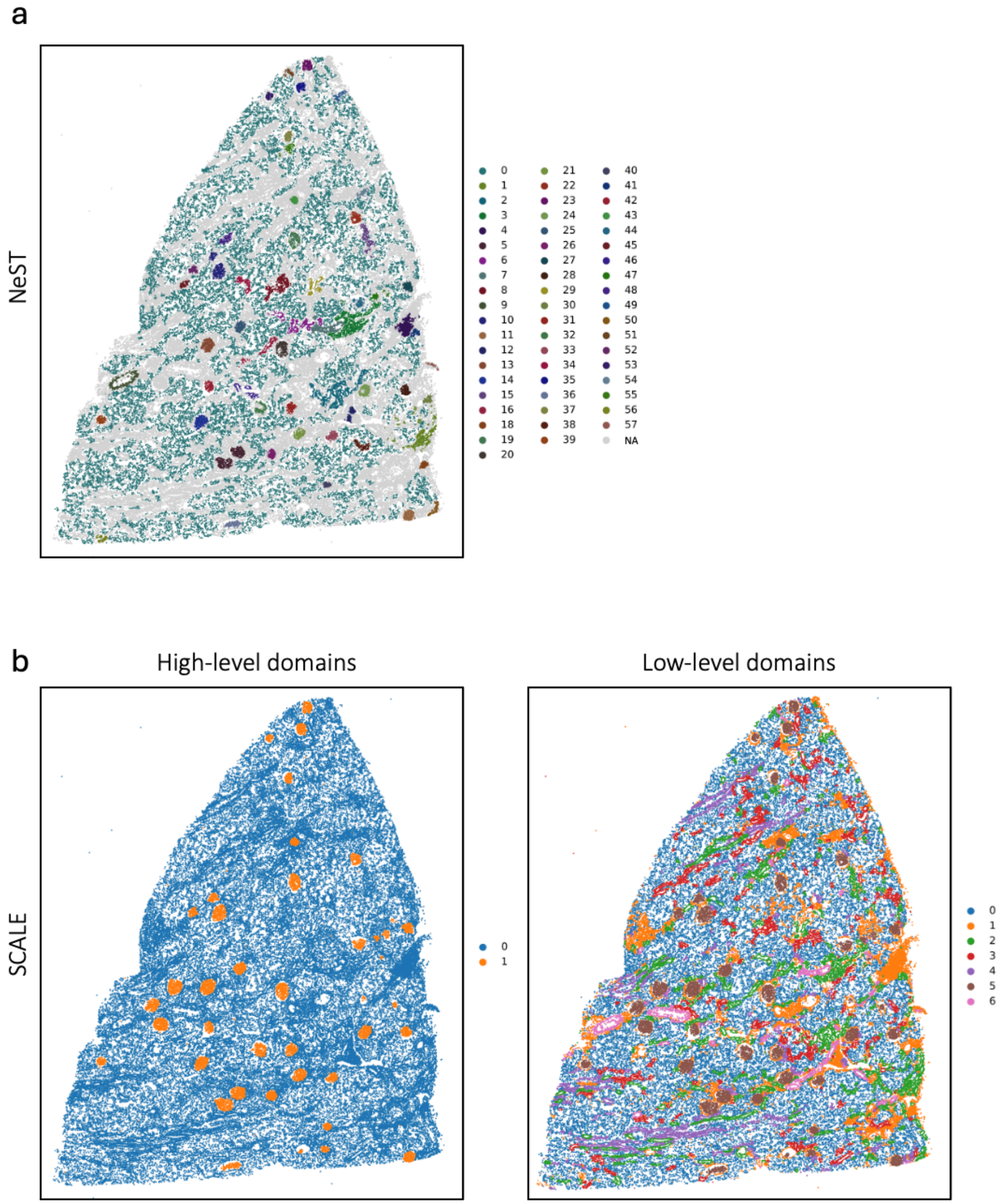


**Supplementary Fig. 5.** Clusters identified for the kidney tissue we used in our study **(a)** NeST and **(b)** SCALE at two scales. NeST identifies 58 clusters and an undefined "NA" group, while SCALE identifies two and seven clusters for each scale, respectively.

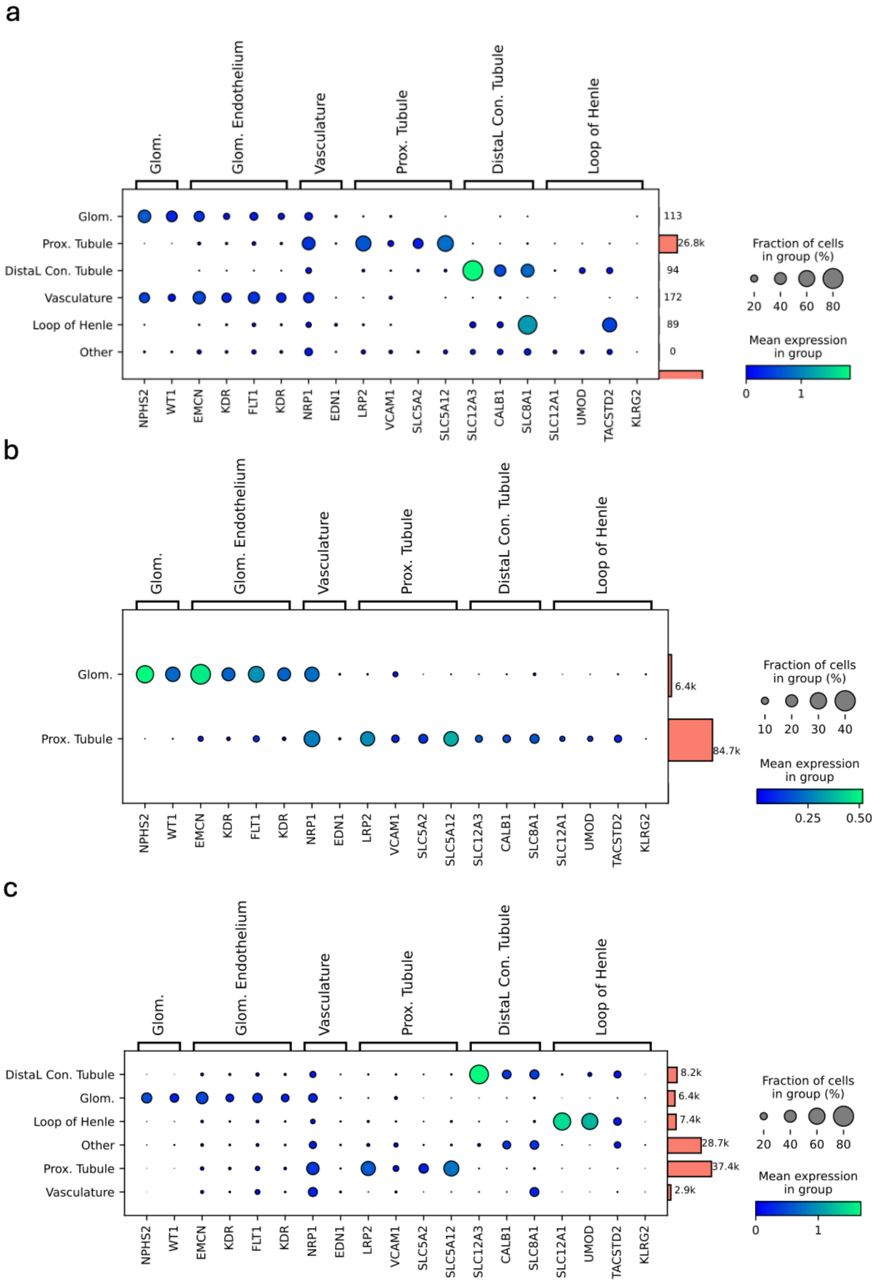


**Supplementary Fig. 6.** Dot plots showing the extent of marker gene expression for each annotated compartment. Dot colors indicate mean-normalized expressions of each marker gene within a compartment and dot sizes indicate the percentage of cells within each compartment expressing the respective marker gene. Clusters were identified using **(a)** NeST, **(b)** SCALE at higher resolution, and **(c)** SCALE at lower resolution. The marginal bar plots show the average mean expression per domain.

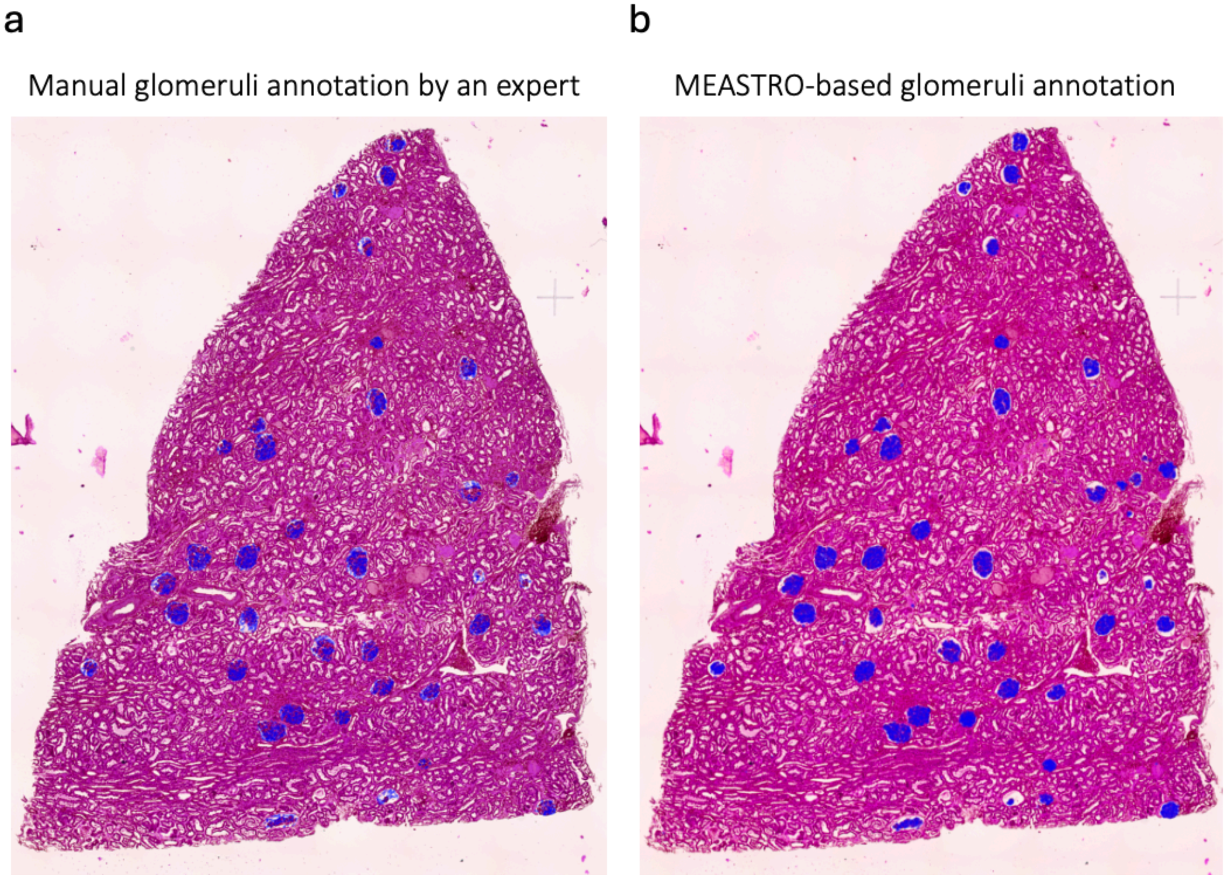


**Supplementary Fig. 7.** Hematoxylin and eosin (H&E)-stained image of kidney tissue showing glomeruli annotations by **(a)** a human expert and **(b)** SCALE. Both sets of annotations are shown in blue. The annotated regions exhibit strong alignment, with SCALE achieving a sensitivity of 100% and a specificity of 88% at the glomerulus level.
